# Abstract spatial, but not body-related, visual information guides bimanual coordination

**DOI:** 10.1101/063404

**Authors:** Janina Brandes, Farhad Rezvani, Tobias Heed

## Abstract

Visual spatial information is paramount in guiding bimanual coordination, but anatomical factors, too, modulate performance in bimanual tasks. Vision conveys not only abstract spatial information, but also informs about body-related aspects such as posture. Here, we asked whether, accordingly, visual information induces body-related, or merely abstract, perceptual-spatial constraints in bimanual movement guidance. Human participants made rhythmic, symmetrical and parallel, bimanual index finger movements with the hands held in the same or different orientations. Performance was more accurate for symmetrical than parallel movements in all postures, but additionally when homologous muscles were concurrently active, such as when parallel movements were performed with differently rather than identically oriented hands. Thus, both perceptual and anatomical constraints were evident. We manipulated visual feedback with a mirror between the hands, replacing the image of the left with that of the right hand and creating the visual impression of bimanual symmetry independent of the right hand’s true movement. Symmetrical mirror feedback impaired parallel, but improved symmetrical bimanual performance compared with regular hand view. Critically, these modulations were independent of hand posture and muscle homology. Thus, vision appears to contribute exclusively to spatial, but not to body-related, anatomical movement coding in the guidance of bimanual coordination.

## Introduction

Whether we type on a keyboard, unscrew a lid, or ride a bike – bimanual coordination is crucial in many of our everyday activities. Therefore, the principles that guide bimanual coordination have received much interest, not least to inform treatment to restore regular bimanual function in clinical settings. Beyond therapeutic considerations, coordinative action can be viewed as an ecologically valid model to understand the principles of movement planning (Oliveira & Ivry, 2008). Accordingly, experiments have studied the factors that constrain bimanual movement execution. A prominent and consistent finding has been that humans can perform symmetrical movements – with symmetry usually defined relative to the sagittal body midline – with higher precision and at higher speeds than parallel movements (Cohen, 1971; Kelso, 1984; Kelso, Scholz, & Schöner, 1986). During symmetrical movements, the two effectors move towards opposite sides of space; for instance, one hand moves to the right while the other concurrently moves to the left. Conversely, parallel movements implicate movements towards the same direction of space; for instance, both hands synchronously move to the left or to the right.

The symmetry bias has been demonstrated across a variety of effectors and movement types, such as finger flexion and extension (Carson & Riek, 1998; Riek, Carson, & Byblow, 1992), finger tapping (Mechsner, Kerzel, Knoblich, & Prinz, 2001), wrist movements (Cohen, 1971), line drawing (Bogaerts, Buekers, Zaal, & Swinnen, 2003), elbow flexion and extension (Spencer & Ivry, 2007), and circling arm movements (Semjen, Summers, & Cattaert, 1995). Given its stability across many qualitatively different movements, symmetry is thought to constitute a general organizing principle of bimanual coordination (Swinnen, 2002). One popular experimental paradigm has been finger abduction and adduction, that is, sideways movements of the two index fingers with the hands held palm down. Participants perform these movements rhythmically, and we therefore refer to this task as "finger oscillations". With the palms down, movement accuracy is high when both fingers are abducted at the same time, resulting in symmetrical finger movements. Accuracy is lower when one finger is abducted while the other one is concurrently adducted, resulting in parallel finger movements (Kelso, 1984).

The mechanisms underlying the symmetry bias have been under debate. Early reports suggested that it originates from anatomical constraints within the motor system, that is, from interactions rooted in muscle synergies caused by hemispheric crosstalk (Cohen, 1971; Kelso, 1984; Riek & Woolley, 2005). Muscle synergies may arise through reciprocal connections between the cortical regions that control homologous muscles of the two body sides and result in preferred activation of homologous limb movements. In this view, symmetrical movements are stable because they involve the same muscles in both limbs, allowing efficient integration of contra- and ipsilateral motor signals. In contrast, parallel finger movements involve different muscles in the two limbs, resulting in reduced stability due to ongoing interference from conflicting ipsi- and contralateral muscle commands (Shea, Buchanan, & Kennedy, 2016).

However, others have suggested that, instead, the symmetry bias originates from interactions rooted in perception (Bingham, 2004; Mechsner et al., 2001). The key finding supporting this proposal was that the symmetry bias prevailed when participants performed oscillatory finger movements with the two hands held in opposite orientations, that is, one palm facing up and the other down. In this situation, symmetrical movements involve non-homologous muscles, whereas parallel movements are achieved through homologous muscles. The persistent advantage of symmetrical over parallel movements despite a reversal of the muscles involved in the bimanual movement is at odds with the idea that muscle synergies alone are responsible for the symmetry bias (Bingham, 2004; Mechsner et al., 2001; Shea et al., 2016).

Several studies have suggested that the previous findings of external vs. anatomical symmetry constraints are not a contradiction, but that both factors jointly influence coordination behavior (Oliveira & Ivry, 2008; Spencer & Ivry, 2007; Swinnen et al., 1998; Temprado, Swinnen, Carson, Tourment, & Laurent, 2003). According to this view, anatomical and external contributions flexibly determine bimanual coordination with their relative weighting depending on context and task demands (Shea et al., 2016). In line with this proposal, we recently observed that the perceptual symmetry bias in the finger oscillation task coexisted with an advantage for using homologous muscles (Heed & Röder, 2014), rather than relying on perceptual coding alone, as had been previously suggested (Mechsner et al., 2001).

Whereas the role of perceptual and anatomical codes has, thus, been firmly established, it is less clear what kind of perceptual information these biases are based on. The prevalent experimental approach has been to contrast vision with posture, and to interpret performance biases induced by vision as evidence for perceptually induced, spatial guidance, and biases induced by posture as evidence for anatomical constraints of movement coordination (Mechsner et al., 2001; Riek & Woolley, 2005). Yet, visual information transports not just abstract spatial information, but also information about the body, presumably to contribute to the construction of a body representation. Indeed, we have found that muscle homology affected bimanual finger oscillations less in congenitally blind than in sighted individuals; this finding suggests that vision may induce not just a spatial bias, but may, in addition, contribute body-related, such as postural and muscle-related, information for motor coordination (Heed & Röder, 2014).

One experimental method to investigate the role of body-related visual information is the use of mirror visual feedback. A mirror is placed along the body midline in the sagittal plane; participants look into the mirror from one side, so that the view of the hand behind the mirror is occluded and replaced by the mirror image of the still visible hand. Thus, although one arm is hidden from view, participants have the impression of seeing both of their hands moving in synchrony (Medina, Khurana, & Coslett, 2015). The strong influence of this visual manipulation on body-related, anatomical aspects is maybe most impressively demonstrated in mirror visual feedback therapy (MVT). MVT is used to treat pathological conditions involving unilateral upper extremity pain and motor dysfunction. The mirror replaces visual feedback of the affected arm with that of the intact arm. Viewing mirrored hand movements of the intact arm has been reported to aid recovery of upper extremity function and/or to alleviate pain in different pathological conditions, including stroke, complex regional pain syndrome, and orthopedic injuries, and can even reduce phantom pain after limb amputation when the mirror image of the remaining hand fills the place of the now missing limb (for reviews see: Deconinck et al., 2014; Ezendam, Bongers, & Jannink, 2009; Moseley, Gallace, & Spence, 2008; Ramachandran & Altschuler, 2009). Thus, in such setups, the visual manipulation of anatomical aspects strongly modulates perception.

Mirror setup can also increase movement coupling between the hands, that is, bimanual symmetrical movements are spatially more similar when mirror visual feedback is available, relative to when only one hand is visible (Franz & Packman, 2004). In the finger oscillation paradigm, mirror feedback can create incongruence between the visually perceived and the truly performed bimanual movement; for instance, during parallel finger movements, mirror feedback feigns symmetrical movement through vision while proprioceptive information signals the true, parallel movement. In this incongruent situation, performance declines compared to regular viewing of the hands and relative to when vision is prevented entirely by closing the eyes (Buckingham & Carey, 2008). In other experimental paradigms, such incongruent visual feedback can even induce phantom sensations, such as tickling or numbness, in healthy participants (Daenen, Roussel, Cras, & Nijs, 2010; Foell, Bekrater-Bodmann, McCabe, & Flor, 2013; McCabe, Haigh, Halligan, & Blake, 2005; Medina et al., 2015).

Thus, a large body of evidence suggests an important role of vision for bimanual coordination, but the specific role of vision for the different aspects to which it can contribute, such as abstract spatial or body-related information, is less clear. One account, the perception-action model put forward by Bingham and colleagues, posits that bimanual coordination performance critically depends on the performer’s ability to perceptually detect the phase relationship between the two limbs, expressed in their relative movement directions (Bingham, 2004; Bingham, Schmidt, & Zaal, 1999; Bingham, Zaal, Shull, & Collins, 2001; Zaal, Bingham, & Schmidt, 2000). Thus, the model specifies visual direction as the aspect of visual information that is relevant for coordination. Difficulty in reliably detecting relative direction presumably leads to maladaptive error detection and correction, which, in turn, impedes performance (Bingham, 2004; Bingham et al., 1999, 2001; Zaal et al., 2000). According to Bingham’s model, bimanual coordination, then, is but a special case of any form of visually driven coordination. In fact, they point out that similar constraints appear to govern coordination of a single limb with either a visual stimulus or the limb of another person (Schmidt, Carello, & Turvey, 1990; Temprado et al., 2003; Wilson, Collins, & Bingham, 2005b; Wimmers, Beek, & van Wieringen, 1992). Accordingly, most experiments that have explored Bingham’s theory have employed paradigms that required unimanual coordination of a limb with moving visual stimuli presented on a display (Snapp-Childs, Wilson, & Bingham, 2011; Wilson, Collins, & Bingham, 2005a; Wilson et al., 2005b; Wilson, Snapp-Childs, & Bingham, 2010). However, this experimental approach implicitly presumes that the brain abstracts from all movement parameters and, in particular, that it dismisses other body-specific, body-related visual information. Yet, the findings that have demonstrated an influence of anatomical in addition to perceptual factors (Heed & Röder, 2014; Spencer & Ivry, 2007; Swinnen et al., 1998; Temprado et al., 2003) suggest that also visual information pertaining to posture and muscles may be of relevance for bimanual coordination.

Here, we used the finger oscillation task as a strictly bimanual paradigm to scrutinize the proposal that bimanual coordination relies predominately on visual direction information, and to integrate the findings from visuo-motor and bimanual coordination that have used different experimental paradigms. The finger oscillation task allowed us to disentangle the three body-related visual aspects that could each potentially be relevant for successful bimanual coordination: first, visual feedback about the spatial direction implied by visual feedback of the performed movement (parallel vs. symmetrical); second, visual feedback about the posture of the hands (same vs. different orientation); and third, visual feedback about the muscles involved in executing the movements (homologous vs. non-homologous).

We conducted the present study to delineate the role of these three aspects of visual information for bimanual coordination. Participants executed oscillatory finger movements that were either parallel or symmetrical relative to the sagittal body midline, with the two hands held either in the same or in different orientations. Participants either viewed their two hands directly, or alternatively viewed their left hand directly and its mirror image at the location in space occupied by the hidden right hand.

## Results

Twenty participants performed the finger oscillation task, that is, they made symmetrical and parallel finger abduction and adduction movements with the index fingers of the two hands with gradually increasing speed (Heed & Röder, 2014; Mechsner et al., 2001). In different blocks, the two hands had either the same orientation with both palms up or down, or different orientations, with one hand facing palm up and the other palm down. This latter manipulation reverses the muscles involved in symmetrical vs. parallel movements: whereas symmetrical movements usually require the use of homologous muscles in the two hands, this muscle configuration is now required for parallel movements. To manipulate visual afferent information, a mirror was placed between the hands in half of the experiment; it hid the right hand, and participants saw the mirror image of the left hand in its place, creating the impression that the currently performed movement was symmetrical, and that both hands had the same posture, independent of the true movement type and hand posture. We tested how the congruence and incongruence of these aspects of visual feedback with the truly performed movement affected the accuracy of bimanual movement coordination.

### Anatomical and external contributions to bimanual coordination

We first tested whether both external and anatomical influences were at all present in our study; the following analyses then focused on which type of visual information modulated these biases. We compared conditions in which correct performance required the use of homologous and non-homologous muscles in the two hands to make symmetrical or parallel movements. We dichotomized movement accuracy by classifying movements as correct when the phase difference of the two fingers deviated by less than 50° from the instructed movement phase in a single movement cycle of abducting and adducting the fingers (180° in external space for symmetrical movements, often referred to as 0° when referring to muscles instead; 0° in external space for parallel movements; Heed & Röder, 2014; Mechsner et al., 2001). If bimanual coordination were solely constrained by anatomical factors, performance should be superior whenever homologous muscles as opposed to non-homologous muscles must be used, regardless of hand posture and movement instruction. Alternatively, if movement coordination were solely constrained by external factors, the symmetry advantage should prevail regardless of whether homologous muscles are involved in the instructed movement. If both anatomical and external factors constrained bimanual coordination, performance in either movement condition should benefit from the use of homologous muscles, in addition to a general advantage of symmetrical over parallel movements.

Whether the instructed movement required the use of homologous muscles depended on the experimental factors movement instruction and hand posture. When both palms had the same orientation, symmetrical movements involved homologous muscles, and parallel movements involved non-homologous muscles. In contrast, when the hands were held in different postures, symmetrical movements involved non-homologous muscles, and parallel movements involved homologous muscles.

Performance declined with increasing movement speed, but more so for parallel than for symmetrical movements, evident in a stronger decline of movement cycles in which the phase difference was classified as correct (i.e., deviating maximally +/-50° from the expected phase difference of 180° for symmetrical, and 0° for parallel movements). In addition, performance was better with the hands in the same than in different postures for symmetrical movements, whereas the opposite performance pattern emerged for parallel movements (Figure 1: left panels; Figure 2). We assessed the statistical significance of these performance differences with a Bayesian model that included parameters that reflected main effects of the experimental manipulations of movement instruction (symmetrical, parallel), hand posture (same, different), and movement speed (dichotomized into slow, fast) and all interactions between them. The posterior distributions of the beta weights that together reflected modulations by anatomical and external spatial coding (β_instruction_, β_posture_, β_speed_, β_instruction_posture_, β_instruction_speed_, β_instruction_posture_speed_) did not span zero, confirming that each factor, as well as their interactions, contributed to bimanual coordination performance (Table 1, Figure 3).

**Figure 1:**
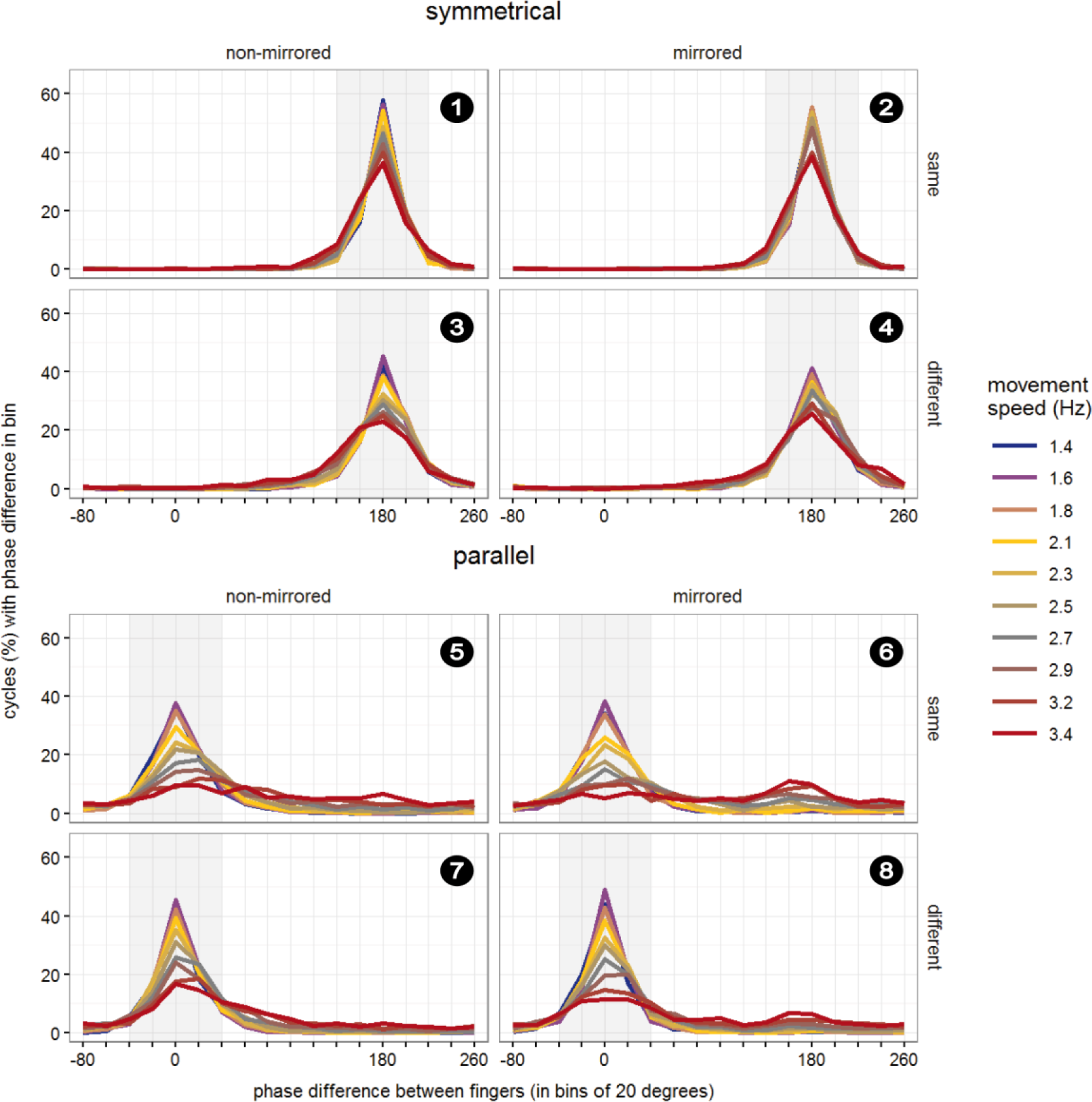
Performance in the finger oscillation task. Relative phase difference was binned in intervals of 20° from −90° to +270° and then divided by the total number of cycles within participants to derive percentage values. Results were averaged across participants, separately for symmetrical (upper panels) and parallel movements (lower panels) at 10 movement speeds. Performance is depicted for non-mirrored (left column) and mirrored (right column) visual feedback conditions, as well as for same (upper panel), and different hand orientations (lower panel). Symmetrical and parallel movements are defined in terms of the horizontal spatial dimension: 180° phase difference indicates moving in perfect symmetry, because one hand is at its leftmost, while the other one is at its rightmost location. In contrast, a 0° phase difference indicates moving perfectly in parallel, because both hands are at their left- and rightmost positions at the same time. Grey shading indicates the range of the phase difference considered as “correct” for statistical analysis (180° +/- 50°: correct symmetrical movement vs. 0° +/-50°: correct parallel movement). Panels are numbered chronologically for integration of results with Figure 2 and Figure 4 (white numbers on black circles).

**Figure 2:**
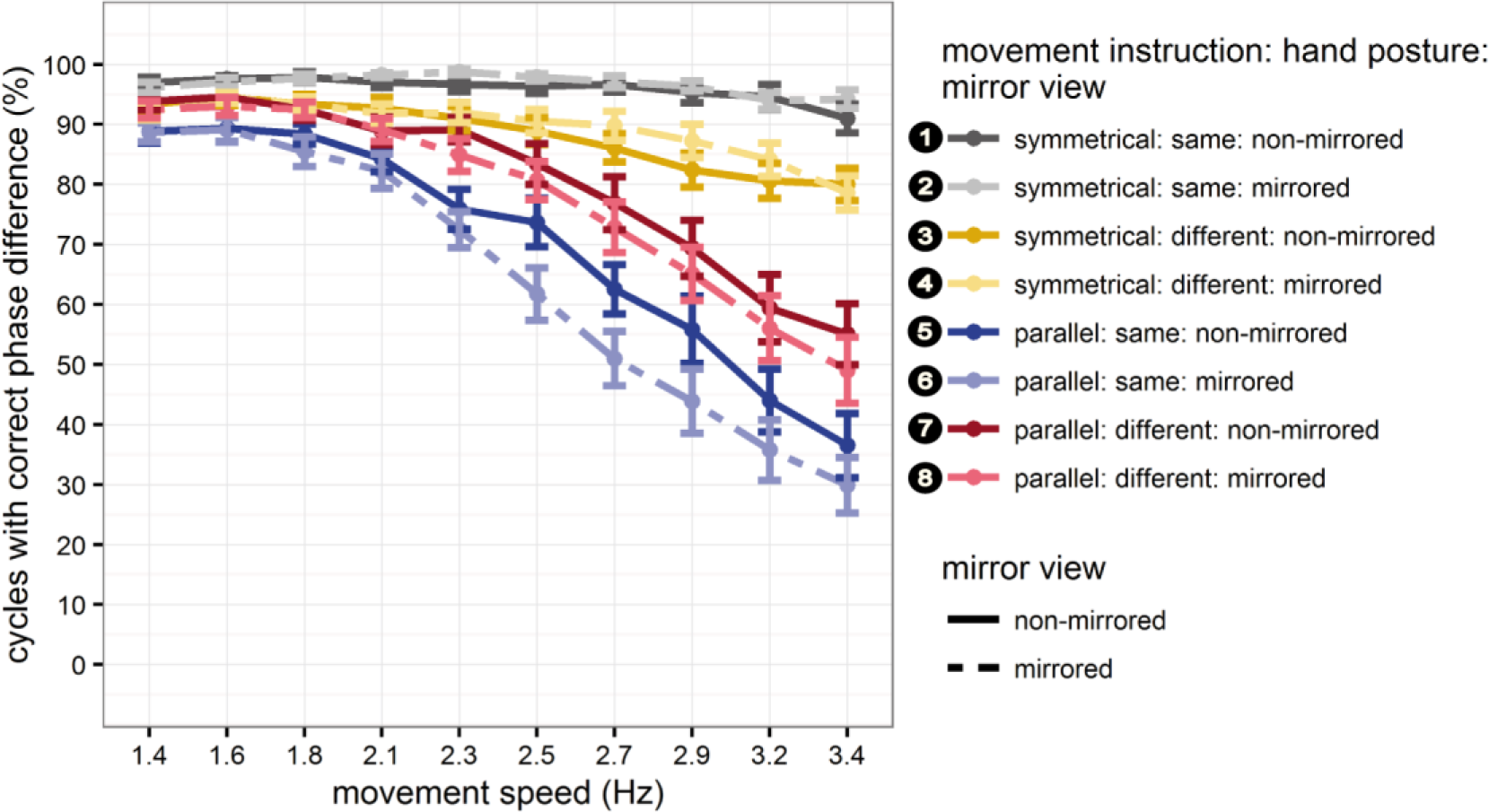
Accuracy in the finger oscillation task. Data points correspond to the grey regions in Figure 1. Percentage of movement cycles with the correct phase difference (+/-50°, as explained in Figure 1) between the two index fingers. Line colors represent the interaction of movement instruction (symmetrical vs. parallel) and hand posture (same vs. different). Dark colors and solid lines represent non-mirrored conditions, and bright colors and dashed lines indicate mirrored feedback conditions. Error bars represent standard errors of the mean. Conditions are numbered in correspondence to Figure 1 and Figure 4 (white numbers on black circles).

**Figure 3:**
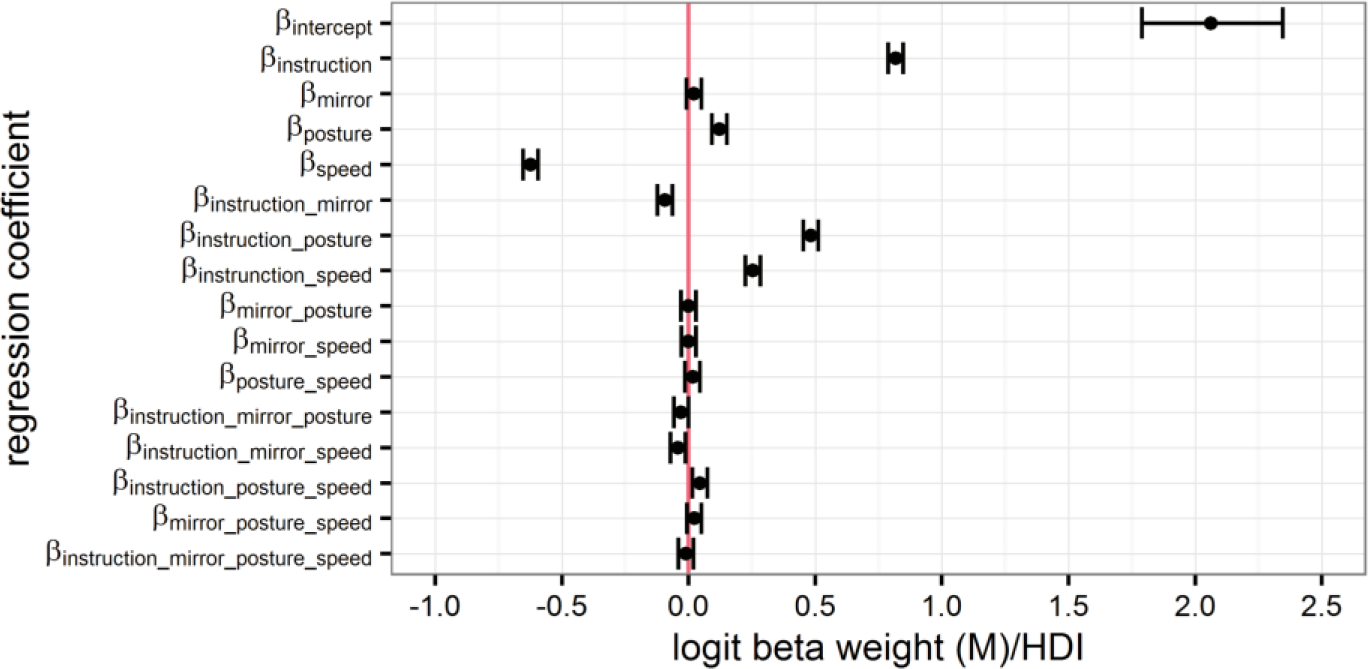
Illustration of the results of the statistical analysis listed in Table 1. Logit mean posterior beta weights of the Bayesian hierarchical logistic regression model. The area between whiskers represents the highest density interval (HDI) of a beta weight’s posterior distribution, as estimated with Markov Chain Monte Carlo (MCMC) sampling. If a beta weight contributes to the prediction of movement accuracy in the finger oscillation task, its HDI does not span zero (depicted as vertical red line).

**Table 1:** Results of the statistical analysis. Logit mean posterior beta weights, their lower (LL) and upper (UL) 95% highest density interval (HDI) limits, and their effective sample size (ESS) of the Bayesian hierarchical logistic regression model, estimated with Markov Chain Monte Carlo (MCMC) sampling. Beta distributions of parameters that were relevant to the tested hypotheses of the study had a minimum ESS of 11,550, ensuring stable and accurate MCMC sampling and chain convergence. Grey shading marks posterior beta weights with an HDI that does not span zero. See Figure 3 for graphical illustration of model results and the text for details on the inferential strategy.

|  | mean | 95% HDI |  | EES |
| --- | --- | --- | --- | --- |
|  |  | LL | UL |  |
| $\beta_{\text{intercept}}$ | 2.06 | 1.79 | 2.35 | 563 |
| $\beta_{\text{instruction}}$ | 0.82 | 0.79 | 0.85 | 12000 |
| $\beta_{\text{mirror}}$ | 0.02 | -0.01 | 0.05 | 12000 |
| $\beta_{\text{posture}}$ | 0.12 | 0.09 | 0.15 | 12000 |
| $\beta_{\text{speed}}$ | -0.62 | -0.65 | -0.59 | 12000 |
| $\beta_{\text{instruction\_mirror}}$ | -0.09 | -0.12 | -0.06 | 12000 |
| $\beta_{\text{instruction\_posture}}$ | 0.48 | 0.45 | 0.51 | 12000 |
| $\beta_{\text{instruction\_speed}}$ | 0.25 | 0.22 | 0.28 | 12000 |
| $\beta_{\text{mirror\_posture}}$ | 0.00 | -0.03 | 0.03 | 12000 |
| $\beta_{\text{mirror\_speed}}$ | 0.00 | -0.03 | 0.03 | 12000 |
| $\beta_{\text{posture\_speed}}$ | 0.02 | -0.01 | 0.05 | 12000 |
| $\beta_{\text{instruction\_mirror\_posture}}$ | -0.03 | -0.06 | 0.00 | 11775 |
| $\beta_{\text{instruction\_mirror\_speed}}$ | -0.04 | -0.07 | -0.01 | 11550 |
| $\beta_{\text{instruction\_posture\_speed}}$ | 0.05 | 0.02 | 0.07 | 12000 |
| $\beta_{\text{mirror\_posture\_speed}}$ | 0.02 | -0.01 | 0.05 | 12000 |
| $\beta_{\text{instruction\_mirror\_posture\_speed}}$ | -0.01 | -0.04 | 0.02 | 12000 |

Hypothesis-driven, direct comparison of the model posterior predictions for conditions that involved homologous vs. non-homologous muscles, separately for symmetrical and parallel movements at slow and fast speeds (parameter: β_instruction_posture_speed_), revealed two key findings. First, the resulting credible difference distributions did not span zero, and the estimated mean performance was larger for symmetrical than parallel movements, both at slow and fast speeds. This result confirmed superior performance of symmetrical over parallel movements independent of hand posture, implying external-spatial contributions to performance. Second, all resulting credible difference distributions were positive, suggesting that performance benefitted from the use of homologous muscles and, thus, indicating that performance was modulated by anatomical factors. These differences were more pronounced at fast than at slow speeds (homologous minus non-homologous conditions: same-different_symmetrical_fast_: M= 2.66 [2.43 2.92]; different-same_parallel_fast_: M= 1.56 [1.43 1.70]; same-different_symmetrical_slow_: M= 2.17 [1.85 2.49]; different-same_parallel_slow_: M= 1.33 [1.13 1.53]).

In sum, these results indicate that bimanual coordination is constrained by external factors, but additionally modulated by anatomical factors, replicating the result of our previous report (Heed & Röder, 2014) in an independent sample and supporting previous accounts of a mixed influence of both in bimanual coordination (Oliveira & Ivry, 2008; Spencer & Ivry, 2007; Swinnen et al., 1998; Temprado et al., 2003).

### Body-related visual information integrated for action

The present study’s main aim was to determine whether, and if so, which specific kind of abstract spatial or body-related visual information constrains movement coordination. Therefore, our experiment was designed to disentangle different kinds of visual feedback: about movement direction, about hand posture, and about the muscles involved in the current action.

Each of these potential influences makes distinct predictions about the pattern of bimanual coordination performance across our experimental factors, and we will briefly introduce each predicted pattern (see Figure 4 for a visual illustration of the three different visual feedback conditions induced by the mirror).

**Figure 4:**
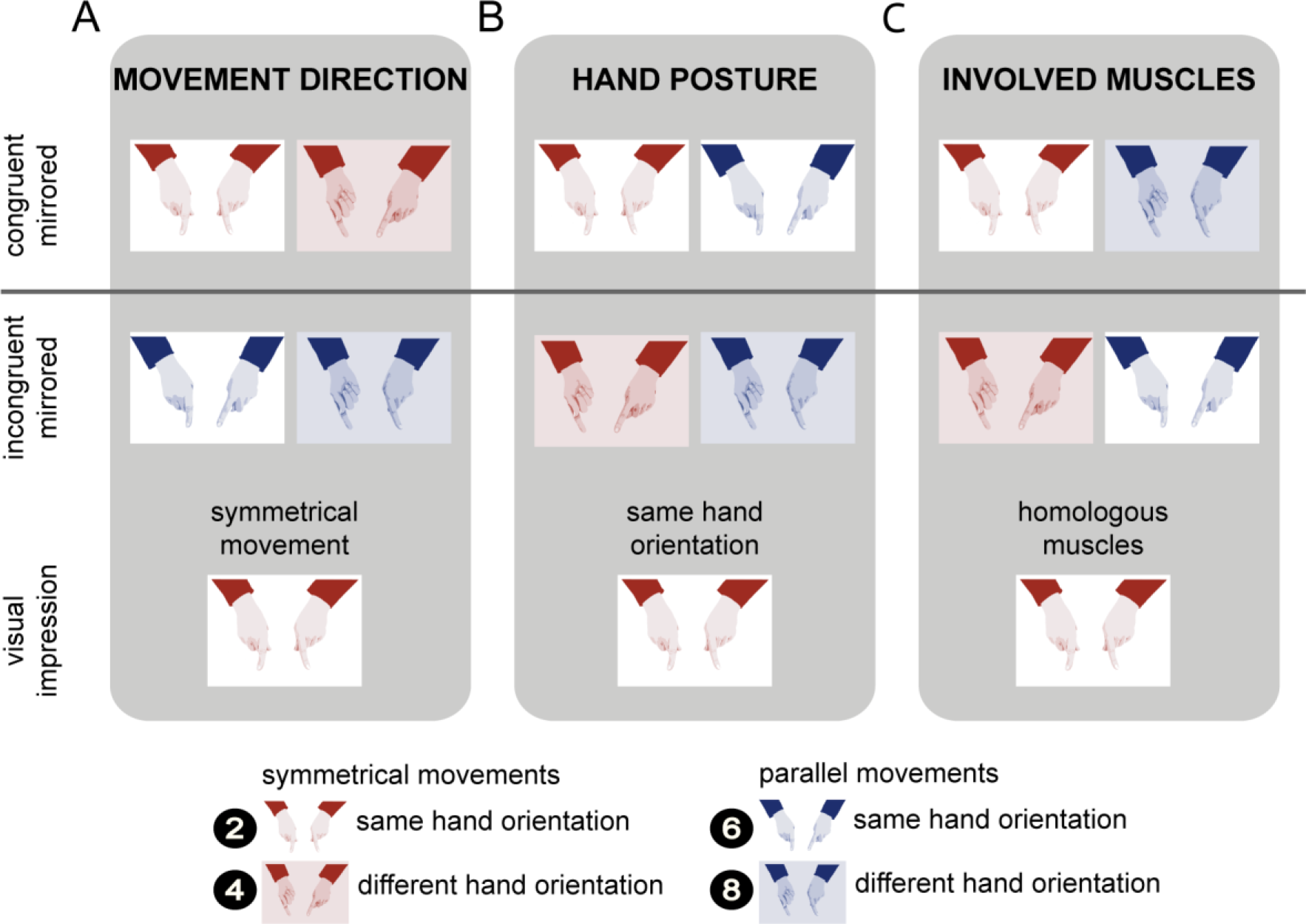
Illustration of the different visual feedback conditions induced by the mirror. Columns structure the mirrored experimental conditions according to visual feedback about movement direction (A), hand posture (B), and involved muscles (C). Rows represent experimental conditions structured according to congruent mirrored, and incongruent mirrored conditions, as well as according to the participants’ visual impression concerning each feedback aspect. Color indicates the movement instruction, with red designating symmetrical movements, and blue parallel movements. Background configuration indicates the hand posture, with no filling designating hands held in the same orientation, and a colored background designating hands held in different orientations. Conditions are numbered in correspondence to Figure 1 and Figure 2 (white numbers on black circles).

#### Visual feedback about movement direction

One potential source of information could be the direction of movement, independent of the further specification of how this movement is achieved, that is, irrespective of posture and involved muscles. In our paradigm, this influence of visual information about movement direction (symmetrical vs. parallel) would be evident in a difference between conditions in which visual and proprioceptive modalities provided congruent versus incongruent information about the type of performed movement (Figure 4A). Without the mirror, visual and proprioceptive information about the executed movement were always congruent (uneven numbered conditions in Figure 1, Figure 2). With the mirror, visual-proprioceptive feedback was incongruent whenever the fingers moved parallel; in these conditions, visual feedback indicated that the fingers were moving symmetrically.

If visual feedback about movement direction were relevant for bimanual coordination, performance in congruent feedback conditions (numbered 2 and 4 in Figure 1, Figure 2, Figure 4A) should be superior to that in conditions with incongruent visual-proprioceptive information (numbered 6 and 8 in Figure 1, Figure 2, Figure 4A). Critically, this difference should be independent of hand posture. Accordingly, congruence of visual-proprioceptive information about movement direction depended on the experimental factors movement instruction and mirror view.

#### Visual feedback about posture

A potential influence of visual information about hand posture would be evident in a difference between conditions with congruent vs. incongruent information about posture from vision and proprioception (Figure 4B). Without the mirror, visual-proprioceptive information about posture was always congruent (uneven numbered conditions in Figure 1, Figure 2). With the mirror, visual-proprioceptive information was incongruent when the two hands had different postures; in these conditions, mirror feedback indicated that the hands had the same orientation. If visual feedback about hand posture were relevant for bimanual coordination, performance should be superior in congruent (numbered 2 and 6 in Figure 1, Figure 2, Figure 4B) over incongruent (numbered 4 and 8 in Figure 1, Figure 2, Figure 4B) visual-proprioceptive posture conditions. Critically, this performance advantage should be independent of movement instruction, that is, of whether executed movements are symmetrical or parallel. Accordingly, congruence of visual and proprioceptive feedback about hand posture depended on the experimental factors mirror view and hand posture.

#### Visual feedback about the involved muscles

A potential influence of visual information about the muscles involved in the current action would be evident in a difference between congruent vs. incongruent visual-proprioceptive information about the currently active muscles (Figure 4C). Without the mirror, visual-proprioceptive information about involved muscles was always congruent (uneven numbered conditions in Figure 1, Figure 2). With the mirror, the combination of movement instruction and hand posture determined whether visual-proprioceptive feedback was congruent or not. Visual-proprioceptive information was, for instance, incongruent when participants made symmetrical movements with differently oriented hands. In this situation, the hands appeared to be oriented in the same posture due to the mirror, and, thus, vision suggested that homologous muscles were used, although truly participants had to use non-homogenous muscles. Further conflict conditions are illustrated in Figure 4C. If visual feedback about muscles were relevant for bimanual coordination, performance in congruent apparent muscle conditions (numbered 2 and 8 Figure 1, Figure 2, Figure 4C) should be superior over incongruent conditions (numbered 4 and 6 in Figure 1, Figure 2, Figure 4C). Accordingly, congruence of visual-proprioceptive feedback about involved muscles depended on the experimental factors movement instruction, mirror view, and hand posture.

### Visual feedback about movement direction is relevant for bimanual coordination

With the mirror present, performance improved for symmetrical movements, but deteriorated for parallel movements, both relative to regular viewing without the mirror. These effects were evident in a gradual decline of the percentage of correctly executed movement cycles with increasing movement speed (Figure 1, Figure 2). For symmetrical movements, this effect was small due to performance near ceiling even at high speeds with the hands held in the same posture. Crucially, the effect of visual feedback varied systematically with movement instruction, but not with hand posture. The posterior distributions of the relevant model beta weights, β_instruction_mirror_ and β_instruction_mirror_speed_, did not span zero, confirming that they contributed to explaining the probability of moving both fingers (Table 1, Figure 3). This result indicates an effect of visual information about movement direction, but not about hand posture and involved muscles.

To further scrutinize this result, we subtracted posterior model predictions in the non-mirrored conditions from those in the mirrored conditions, separately for symmetrical and parallel movements at slow and fast speeds (parameter: β_instruction_mirror_speed_). The credible difference distributions are displayed in Figure 5. Performance deteriorated during parallel movements in mirror as compared to non-mirrored conditions, as evident in the negative distribution of credible differences at both slow and fast speeds, all of which did not span zero. In contrast, performance improved during symmetrical movements in mirrored relative to non-mirrored conditions, as evident in the positive distribution of credible differences at fast speeds, which again did not span zero. This performance improvement was not evident at low speeds, presumably because performance was more similar overall during slow movements, in line with previous reports (Figure 2).

**Figure 5:**
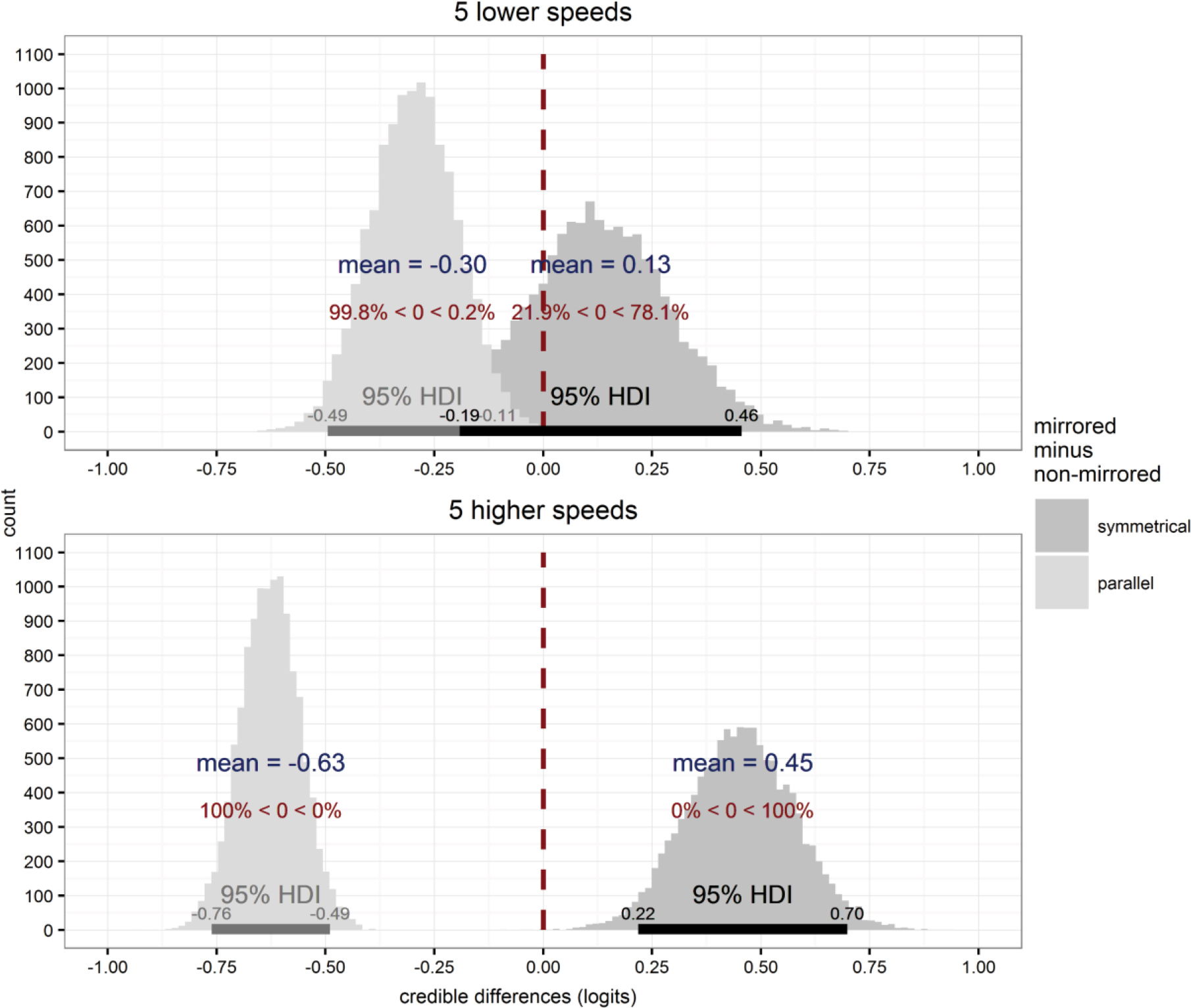
Illustration of credible difference distributions of the parameter βinstruction_mirror_speed estimated within the Bayesian hierarchical logistic regression model. Mirrored and non-mirrored visual feedback conditions are contrasted separately for symmetrical and parallel movements (dark and light grey) across slow and fast movement speeds (top and bottom row). Red inscriptions per distribution indicate the percentage of distributions’ samples falling below and above zero. Horizontal bars indicate 95% highest density interval (HDI) limits. Credible difference distributions indicated that visual feedback about movement direction influenced bimanual coordination. At low speeds, performance deteriorated for parallel movements when the mirror was present (light distribution in upper panel). In contrast, no reliable change was evident for symmetrical movements, evident in that the darker distribution in the upper panel includes zero. At high speeds, too, performance deteriorated for parallel movements when the mirror was present (light distribution in the lower panel), but improved for symmetrical movements with the mirror present as compared to regular viewing (dark distribution in the lower panel).

### Visual information about hand posture and involved muscles are irrelevant for bimanual coordination

To further test whether, indeed, coordination relied solely on visual direction information, we directly examined the parameter estimates relevant for the potential alternatives, namely, hand posture and involved muscles.

For hand posture, the posterior distributions of the model beta weights β_mirror_posture_, and β_mirror_posture_speed_ spanned zero, suggesting that this experimental factor did not contribute to explaining the probability of moving correctly (Table 1, Figure 3). Thus, statistical analysis did not provide any evidence that visual information about hand posture constrained movement coordination in the present experiment.

An effect of visual information about involved muscles would be evident in the interaction of the experimental factors movement instruction, mirror view, and hand posture (Figure 4C). Note that a modulation of visual information about involved muscles would thus encompass the same factors that also indicate a modulation of visual information about movement direction, namely movement instruction and mirror view, but would warrant an additional modulation by hand posture. The posterior distributions of the corresponding model beta weight β_instruction_mirror_posture_ just barely excluded zero (Table 1, Figure 3). Nonetheless, we followed up on this finding by subtracting posterior model predictions for incongruent from congruent mirror conditions, separately for symmetrical and parallel movements. The distributions of credible differences were positive and did not span zero, indicating that performance in congruent feedback conditions was superior to performance in incongruent conditions, as would be predicted if visual information about involved muscles were relevant for coordination (congruent minus incongruent conditions: same-different_symmetrical_mirrored_: M= 2.53 [2.23 2.83]; different-same_parallel_mirrored_: M= 1.56 [1.39 1.72]).

We further reasoned that, if visual feedback about the involved muscles indeed determined coordination, performance in congruent mirror conditions should be indistinguishable from performance in corresponding conditions without mirror, because in both cases, visual and proprioceptive feedback unanimously indicate that corresponding muscles are used. Additionally, along with altering visual feedback concerning muscle identity, the mirror manipulation presumably affected visual feedback concerning the relative timing of bimanual muscle activation. With regular visual feedback of the hands, the dominant hand has been observed to lead the non-dominant hand by about 25 ms in bimanual coordination tasks (Semjen et al., 1995). Correspondingly, mirrored feedback about the timing of muscle activation would not correspond exactly to its actual timing, given the slight lag of the non-dominant hand. Therefore, we predicted that performance in congruent mirrored conditions should be worse than in congruent non-mirrored conditions if visual information concerning involved muscles determined coordination. To test this prediction, we subtracted posterior model predictions for congruent non-mirrored from congruent mirror conditions, separately for symmetrical and parallel movements. Note that a differential effect of mirror view depending on movement instruction cannot be accounted for by a visual effect of involved muscles, as both conditions are identical concerning muscle information. If nonetheless the effect of mirror view depends on the movement instruction, this would further corroborate the effect of visual movement direction, as parallel and symmetrical movements differ concerning this aspect.

The effect of mirror view indeed differed according to the movement instruction. Performance improved with mirrored feedback, relative to non-mirrored conditions, when moving symmetrically (mirrored-non-mirrored_symmetrical_same_: M= 0.41 [0.06 0.76]). The opposite pattern was evident when moving in parallel, that is, mirrored visual feedback was detrimental to performance (mirrored-non-mirrored_parallel_different_: M= -0.35 [-0.52 -0.16]).

Contrary to the comparison of congruent vs. incongruent mirrored conditions concerning involved muscles, the comparison of congruent mirrored with congruent non-mirrored conditions, thus, did not support the notion that visual feedback about the involved muscles constrains bimanual coordination. Instead, the credible, but differential effect of mirrored visual feedback on performance depended on the movement instruction and corroborates that visual movement direction affected coordination performance.

### Temporal aspects of visual feedback concerning movement direction

The performance improvement during the viewing of mirrored symmetrical feedback struck us as surprising, as one might expect that the perception of non-veridical visual movement timing feedback would be detrimental to, rather than supportive of, the production of coordinated movement. The present finding led us to speculate that the temporal synchrony of visual feedback in the mirrored condition may actually lead to a decrease of the true lag between the dominant and non-dominant hands in our experiment, potentially marking a mechanism by which the mirror-induced performance improvements observed here may be explained.

When movement direction was visually and proprioceptively congruent, performance was better in mirrored than non-mirrored conditions; this difference was small, but associated with a credible difference parameter estimate in our model. Performance of symmetrical movements was generally near ceiling, so that even substantial differences on the logit scale translate to very small differences in performance measured as percentage correct. Accordingly, the 0.45 improvement on the logit scale translates to only a 0.3% percentage correct improvement at high movement speeds (beta weight in the model: β_instruction_mirror_speed_). Conversely, smaller differences on the logit scale in other conditions were much more clearly evident on the percentage correct scale. The performance improvement with mirrored relative to non-mirrored feedback (beta weight in the model: β_instruction_mirror_posture_) and hands held in different orientations was estimated at 2.3% (logit: 0.19; baseline performance level: 85.2%, logit: 1.75), as compared to a 1.0% (logit: 0.26, base performance level: 95.5%, logit: 3.05) improvement with hands held in the same orientation. Nonetheless we are hesitant to capitalize on this result, as the beta weight including posture (beta weight in the model: β_instruction_mirror_posture_) just barely excluded zero and the performance decline when performing parallel movements with the mirror present relative to non-mirrored visual feedback, was larger (13.7%; 0.63 logits; beta weight in the model: β_instruction_mirror_speed_).

## Discussion

The present study aimed at specifying anatomical and external-spatial contributions to bimanual coordination performance. Previous findings, mainly from experiments requiring the coordination of limb movements with visual cues, have led to a theoretical account of bimanual coordination, and motor coordination more generally, that stresses the relevance of the perceivability of phase synchrony implied in visual direction information (Bingham, 2004; Bingham et al., 1999, 2001; Zaal et al., 2000). In contrast, findings from some bimanual coordination paradigms have stressed the importance also of anatomical factors such as the muscles involved in a particular bimanual movement, suggesting that visual information about factors other than solely movement direction may play a role in coordinative behavior of the limbs (Heed & Röder, 2014; Swinnen et al., 1998; Temprado et al., 2003). We exploited the well-known bias towards symmetrical over parallel finger movements to delineate different potential sources of visual modulation by introducing a mirror through which participants saw the reflection of one hand projected onto the location of the hidden, other hand. Our study revealed three key results. First, anatomical factors modulated bimanual coordination. Specifically, participants performed better when bimanual movements required the concurrent activation of homologous rather than non-homologous muscles. Second, external spatial factors, too, modulated bimanual coordination. An advantage of symmetrical movements prevailed regardless of hand posture, and, thus, irrespective of whether homologous muscles had to be activated. Third, of the three kinds of visual information manipulated in the present study – movement direction, hand posture, and the muscles involved in the performed movements –, only movement direction information modulated bimanual performance. In contrast, visual information pertaining to hand posture appeared to be irrelevant for coordination performance, and there was only weak evidence that visual information pertaining to the muscles involved in the current movement may play a role in coordination performance.

In line with the specific modulation by visual direction information we observed in the present experiment, previous studies have demonstrated that visual directional cues are relevant for bimanual coordination. For instance, most coordination tasks result in inherently stable performance only when the bimanual phase patterns are symmetrical or parallel, but not for intermediate phase differences (Kelso, 1984). Yet, participants can execute such out-of-phase movements if their movement is yoked to concurrent symmetrical or parallel visual information while the hands are hidden from view. For instance, human participants can execute four circular hand movements with one hand, and concurrently five with the other only if these movements are translated into equally fast visual circular movements (Mechsner et al., 2001; see Tomatsu & Ohtsuki, 2005 for a similar finding). Furthermore, performance of orthogonal bimanual movements, such as one hand moving up and down, while the other hand moves to the left and right, improves if visual feedback is given in one plane, that is, as if both hands were moving up and/or down (Bogaerts et al., 2003). These studies suggest that performance of less stable coordination patterns improves if directional visual feedback indicates that an inherently stable coordination pattern, that is, symmetrical or parallel movement, is performed.

Bimanual movements can also be stable when visual feedback is not symmetrical or parallel, but if, instead, movement paths of both hands can be visually perceived as forming a common, coherent shape (Franz, Zelaznik, Swinnen, & Walter, 2001). In a similar vein, participants can execute polyrhythmic two-hand movements when guided by visual displays that integrate directional information of the two hands into one common visual signal (Shea et al., 2016). These so-called Lissajous displays integrate the position of the two hands into a single point on the display by mapping the movement of each limb onto one axis. Performance in this setup is best if the display shows both the visual target pattern and a cursor that indicates the current (transformed) limb position (Kovacs, Buchanan, & Shea, 2008, 2009, 2010; Kovacs & Shea, 2011). Performance declines rapidly if the display is turned off, suggesting that the integration of the immediate visual direction information about the to-be-performed coordination pattern is a prerequisite for its execution (Kovacs et al., 2008; Kovacs & Shea, 2011).

Kovacs and colleagues (2010) have interpreted these findings as empirical support of the perception-action model proposed by Bingham and colleagues, which capitalizes on visual direction information as the cardinal factor for successful bimanual coordination (Bingham, 2004; Bingham et al., 1999, 2001; Zaal et al., 2000). Visual conditions such as those created by the above-mentioned experimental setups then presumably aid error detection, because they facilitate the perceivability of relative movement direction (Kovacs et al., 2009, 2010). In line with the idea of visual movement direction driving coordinative behavior, typical coordination phenomena, such as the advantage of symmetrical over parallel movements, persist even if movements are coordinated only visually. This is the case, for instance, when two people must coordinate their movements (Schmidt et al., 1990; Temprado et al., 2003) and when participants must coordinate their movement with moving visual stimuli on a display (Wilson et al., 2005b; Wimmers et al., 1992). Using such a visual coordination paradigm, it has been demonstrated for example that training participants abilities’ to detect relative movement direction, improves coordination performance with a moving visual stimulus on a display (Wilson et al., 2010). In a similar vein, perceptual detection of relative phase has been shown to be largely unaffected by alternative candidate movement parameters, such as frequency and speed, thus further scrutinizing the importance of relative movement direction for the perceivability of relative phase (Wilson & Bingham, 2008). In light of these results, it has been suggested that bimanual coordination is but a special case of any form of visually driven coordination and as such similarly relies on the perceptual ability to detect relative phase from movement direction. Crucially, this conclusion presumes that the brain abstracts movement direction and dismisses all other body-specific visual information. We provide direct experimental evidence for this assumption here, using a strictly bimanual paradigm and thus bridging the gap between findings from visuo-motor and bimanual coordination that have used different experimental approaches.

Collectively, then, these results stress the importance of visual movement direction for bimanual coordination and provide a comprehensive account for the dominant role of visual direction information we observed in the present study. In contrast, a general degeneration of vision does not impair performance (Buckingham & Carey, 2008; Mechsner et al., 2001; Swinnen, Lee, Verschueren, Serrien, & Bogaerds, 1997), or, leads to only a minor destabilization (Salesse, Oullier, & Temprado, 2005).

Similarly, visual augmentation by marking fingers that have to move together to produce symmetric or parallel tapping patterns does not affect performance (Mechsner, 2004). Moreover, previous studies have suggested that movement execution is modulated by the level of abstraction of visual effector feedback (Brand et al., 2016; Veilleux & Proteau, 2010). Our study did not abstract visual direction information, but, through the mirror setup, provided participants with visual feedback that appeared to reflect the real hands. This experimental situation, thus, more closely resembles the true visual feedback of everyday situations, in which we usually have full vision of our effectors (Holmes & Spence, 2005). Our results show that the brain indeed abstracts movement direction from body-related visual feedback during bimanual coordination, while discarding visual information regarding hand orientation, as well as involved muscles, and thus validates a generalization of the findings obtained with more abstract feedback situations, such as cursors on a screen, to realistic feedback situations.

It is under debate whether continuous, rhythmic movements and short, goal-directed movements rely on similar brain mechanisms. The role of visual information has been investigated in the context of bimanual goal-directed movement (Reichenbach, Franklin, Zatka-Haas, & Diedrichsen, 2014; C. Weigelt & Cardoso de Oliveira, 2002; M. Weigelt, Rieger, Mechsner, & Prinz, 2007) and especially in the context of unimanual goal-directed movement (e.g., Wolpert, Ghahramani, & Jordan, 1995). In these studies, visual information about effector position affected performance, in line with the requirement of integrating target location with current limb position (Kalaska, Scott, Cisek, & Sergio, 1997; Saunders & Knill, 2003). For instance, visual information about the limb can dominate proprioceptive position, information a phenomenon termed ‘visual capture’ (Hay, Pick, & Ikeda, 1965; Holmes, Crozier, & Spence, 2004). Furthermore, specific resources appear to be devoted to monitoring hand position during goal-directed movement (Reichenbach et al., 2014). The relative contribution of – usually redundant – visual and proprioceptive signals to movement planning depends on the reliability of each informational source (Ernst & Banks, 2002; McGuire & Sabes, 2009; Morgan, DeAngelis, & Angelaki, 2008; Sober & Sabes, 2003; van Beers, Sittig, & Denier van der Gon, 1998, 1999), and the relative weighting of visual and proprioceptive signals differs according to the stage in motor planning (Sarlegna et al., 2003; Sober & Sabes, 2003). Visual information appears to be most important when inferring external spatial movement parameters, whereas primarily proprioceptive feedback is used when inferring muscular-based, position-related information, as is necessary to translate a motor plan into body-or hand-centered coordinates for execution (Sarlegna et al., 2003; Sarlegna & Sainburg, 2009; Sober & Sabes, 2003).

To relate the present study to these findings from studies on goal-related movement, one can conceptualize the present repetitive finger oscillation task in an analogous framework. Here, visual direction information outweighed proprioceptive and motor signals to guide continuous bimanual coordination, in line with the finding goal-directed movements primarily rely on visual information when external spatial movement parameters must be inferred. In contrast, visual information about hand posture and involved muscles did not affect performance, suggesting that proprioceptive information outweighed visual feedback for these properties in the present task. This pattern is in line with the prominent role of proprioceptive signals when muscular-based, position-related information must be derived for goal-directed movement to translate a motor plan into body- or hand-centered coordinates for movement execution. However, the repetitive nature of the present bimanual task prohibits formally distinguishing between planning and execution stages of the movements, and, thus, makes it difficult to draw firm conclusions about the potential overlap regarding the processing principles of goal-directed, unimanual and continuous, bimanual movements.

In the present task, mirrored visual movement information was always integrated for bimanual coordination, but the behavioral consequences of integration depended on whether visual movement information was congruent or incongruent with proprioceptive and motor signals. This result pattern seems to be at odds with previous studies that reported that integration of mirrored visual feedback scaled with the degree of congruency of visual and proprioceptive movement information (Bultitude, Juravle, & Spence, 2016; Holmes, Snijders, & Spence, 2006; Medina et al., 2015). In these studies, synchronous movements led to reliance primarily on visual information, whereas asynchronous movements led to reliance primarily on proprioceptive information. Notably, the dependent measures marking integration of visual information in these studies – gap detection at, or pointing movements with, the hidden hand – were acquired after bimanual movements with mirrored visual feedback had been performed for some time. Thus, the dependent measures were unimanual and as such not indicative of visual contributions to bimanual coordination performance. Furthermore, both measures might differ considerably with regard to the reliability and relevance assigned to bimanual visual information, as compared to continuous bimanual coordination performance assessed in the present task.

Incongruence of movement-related visual, proprioceptive, and motor information led to a performance decline of bimanual coordination in our study. This result is in line with reports of MVT suggesting that incongruent sensory feedback induces phantom sensations, such as tickling and numbness, in healthy participants (Daenen et al., 2010; Foell et al., 2013; McCabe et al., 2005; Medina et al., 2015). In contrast, congruence of mirrored visual, proprioceptive, and motor information led to a performance improvement, possibly because the mirrored movement information during symmetrical movements provided optimized visual feedback about the temporal aspects of bimanual movements. These findings bear relevance on clinical applications of the mirror manipulation. So far, few standardized MVT treatment protocols exist, and those that do have specified that movements should be bilateral and performed in synchrony, but have not stressed that they should be symmetrical as well (Grünert-Plüss, Hufschmid, Santschi, & Grünert, 2008; McCabe, 2011). It has even been suggested that the “[…] actual manner of movement appears not to matter as long as it is bilateral and synchronized” (p.175 McCabe, 2011). Additionally, it has been suggested that therapeutic aids should be used unilaterally using the healthy arm in front of the mirror (Grünert-Plüss et al., 2008). These and similar instructions possibly produce incongruence of proprioceptive and visual movement direction, which might produce undesired effects and explain why scientific evidence in favor of MVT as a tool to aid bimanual function is still scarce to date. Consequently, the selective performance benefit of mirrored symmetrical movements and the detrimental effect of incongruent visual movement information for bimanual coordination we report here suggest that applications of MVT should stringently ensure that congruent, symmetrical movements are performed, and further imply that unimanual mirrored handling of therapeutic aids may be disadvantageous to the facilitation of bimanual coordination.

In conclusion, bimanual coordination is guided both by anatomical, muscle-based constraints, as well as by perceptually based, visual constraints. For the latter, information about direction appears to play a key role, whereas effects of posture and muscle homology appear to be mediated only through non-visual channels, and visual cues pertaining to these aspects did not further modulate performance. These results integrate well with current models of bimanual control and goal-directed movement that posit a guiding role of abstract visual direction information for movement planning and execution.

## Methods

We report how we determined sample size, all experimental manipulations, all exclusions of data, and all evaluated measures of the study. Data and analysis scripts are available online (see https://osf.io/g8jrt/).

### Participants

Previous studies have typically reported significant results pertaining to posture in the finger oscillation task with N<10 (e.g., Heed & Röder, 2014; Mechsner et al., 2001). Here, we defined, in advance, a target sample size of 20 participants because we expected that mirror-induced effects would be smaller than posture effects, requiring a larger number of participants for statistical power. Data were acquired from 23 participants, because the data of 3 participants had to be excluded from analysis (see below). None of the participants had participated in our earlier study (Heed & Röder, 2014). All participants were students of the University of Hamburg. They were right-handed according to self-report (mean laterality quotient: 80.4, range: 50-100; Oldfield, 1971), and had normal or corrected-to-normal vision and did not report any neurological disorders, movement restrictions, or tactile sensitivity problems. They provided written informed consent and received course credit for their participation. The experiment was approved by the ethics committee of the German Psychological Society (DGPs). Two participants aborted the first experimental session after a few trials, because they were unable to perform the bimanual coordination task. Data of a third participant was excluded because movements were accidentally instructed incorrectly. The final sample thus consisted of 20 students, 15 of them female, mean age 23.6 years (range: 20-32 years).

### Experimental design

The experiment was designed based on the studies by Mechsner and colleagues (2001) and Heed and Röder (2014). Figure 6 illustrates the setup and the experimental conditions. Participants performed a finger oscillation task; they executed adduction and abduction movements, that is, right-left movements, with the two index fingers. Instructed movements were either symmetrical, that is, the index fingers moved in- or outwards at the same time, or parallel, that is, fingers moved to the right or left side in space at the same time (see Figure 6B). There were two viewing conditions: non-mirrored and mirrored (see Figure 6A). In the non-mirrored conditions, participants viewed both hands directly and, thus, received regular visual feedback. In the mirrored conditions, a mirror blocked the view of the right hand, so that participants saw the mirror image of the left hand in place of their real right hand; however, this manipulation gives rise to the subjective impression of seeing both hands just like in the non-mirrored condition. The hands were either held in the same (both palms up or down) or in different hand orientations (right palm up, left palm down, or vice versa; Figure 6C).

**Figure 6:**
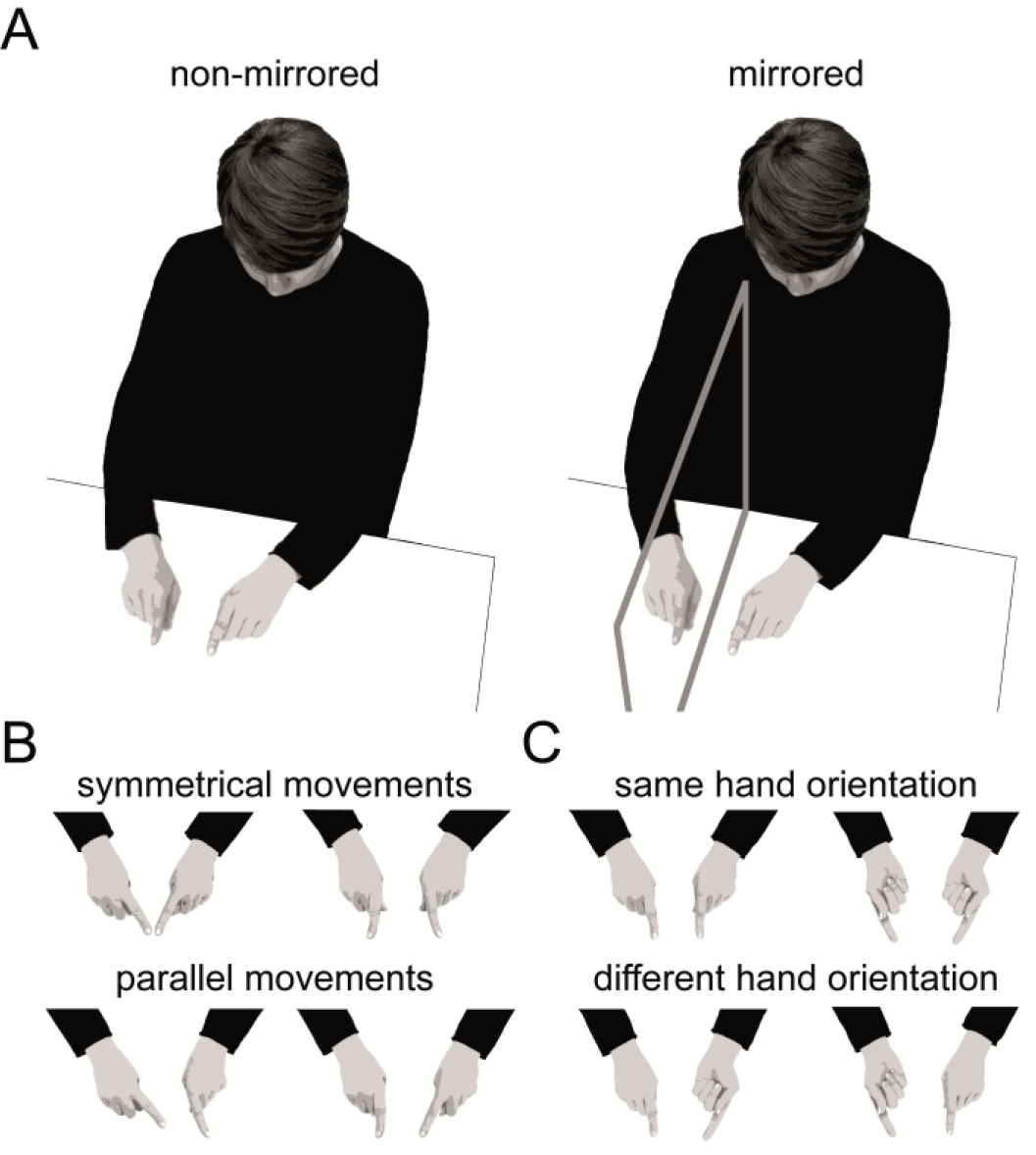
Illustration of the finger oscillation task. Participants performed adduction and abduction movements with the index fingers of both hands. A. Participants either viewed their hands directly, or looked into a mirror, so that they saw their left hand, and the left hand’s mirror image at the location of the right hand. B. For symmetrical movements (upper row), participants concurrently moved both fingers in- and outwards. For parallel movements (lower row), participants concurrently moved the two fingers to the left and right in space. C. Hands were held either in same (upper row) or in different orientations (lower row).

The experiment comprised four experimental factors. The factors movement instruction (symmetrical vs. parallel), mirror view (non-mirrored vs. mirrored), and hand posture (both palms down vs. both palms up vs. left palm up and right palm down vs. right palm up and left palm down) were varied block-wise in randomized order. The factor speed (10 discrete speeds from 1.4 to 3.4 Hz) was varied within trials. Whereas participants are usually able to perform symmetrical and parallel movements (almost) equally well at low speeds, their performance regularly declines markedly for parallel, but not symmetrical, movements at high speeds (Kelso, 1984). During a trial, each speed level was maintained for 5 beats, resulting in 50 beats per trial, resulting in a trial duration of about 22 seconds. Each of the 16 combinations of the factors instruction, mirror view, and hand posture was presented 4 times across two sessions held on separate days.

### Materials and apparatus

Participants sat at a table with both hands resting comfortably in front of the body. Finger movements were tracked with a camera-based motion tracker (Visualeyez II VZ4000v PTI; Phoenix Technologies) using infrared markers sampled at 100 Hz. Four markers were attached to each index finger, one on the finger nail, one opposite the nail on the fingertip, and one on each side between nail and tip. As a result, at least one marker per hand was visible during movement execution in all postures. Movements were instructed by metronome-like sounds presented through two loudspeakers positioned in front of the participant. Experimental protocols were controlled via MATLAB (version 7.14, The Mathworks).

### Procedure

In each trial, participants rhythmically moved both outstretched index fingers to the metronome sounds. Participants were instructed to complete a full movement cycle per beat, that is, move both fingers at the same time in- and outwards when moving symmetrically, or, move both fingers at the same time to the left and right in space when moving in parallel.

Instructions stressed that participants should execute movements as correctly as possible, but could change to a more comfortable movement pattern if they were unable to maintain the instructed movement pattern (Lee, Blandin, & Proteau, 1996). Participants had to look at both hands (both real or left real/right mirrored) throughout the experiment. They rested and stretched after every 2 trials.

### Data selection and trajectory analysis

Two trials from one participant were excluded because the hand position on the table had accidentally been instructed incorrectly. Two trials were excluded because a participant had partially closed his/her eyes to ease performance. We analyzed the left-right component of finger movement trajectories.

Within trials, occasional missing data were interpolated (e.g., if a marker was temporally non-visible), trajectories smoothed with a low-pass filter (first-order Butterworth filter at 7.5 Hz), and normalized by demeaning.

Individual movement cycles were then identified as the interval between a consecutive maximum and minimum of the right finger’s trajectory. A sine wave was fitted to the trajectory of this interval for each finger (see Y.Q. Chen, 2003, http://www.mathworks.com/matlabcentral/fileexchange/3730-sinefit). See Figure S1 in the supplemental results for an illustration of the sine wave fit to the raw data. The relative phase of the two fingers was determined as the phase difference of the two fitted sine curves. For symmetrical movements the phase difference should be 180°, because one finger is at its rightmost position when the other is at its leftmost position. For parallel movements the phase difference should be 0°, because both fingers move in synchrony to the left and right in space.

The final data set consisted of 62,536 movement cycles from 20 participants with an average of 39 movement cycles per condition and participant (range: 25-46). The reasons for the variability of the number of movements are that participants sometimes paused or made unidentifiably small movements, especially at high speeds; furthermore, participants were sometimes off-beat and then executed fewer movement cycles than instructed.

### Statistical inference: Bayesian hierarchical logistic regression

In Bayesian statistical analysis approaches, credibility is reallocated across candidate parameter values, such as slopes indicating the effect of a certain experimental factor for example, as data, also called ‘evidence’, is cumulatively taken into account (Kruschke, 2015). These candidate parameters values are given a certain a priori credibility, called the ‘prior’, which is typically noncommittal. The result of Bayesian model estimation then is a posterior distribution of jointly credible parameter values, given the evidence and the prior belief that certain values are more, less, or equally, likely (Kruschke, Aguinis, & Joo, 2012). Conveniently, the resulting posterior distribution is directly indicative of where in parameter space the true value is most likely to be.

For statistical inference, we dichotomized the phase difference of the two fingers into correct (1) and incorrect (0). To this end, the relative location of the two fingers during a movement cycle was compared to the expected relative difference in each condition (+/- 50° around °0 and 180° for parallel and symmetrical movements, respectively, see Mechsner et al., 2001; Heed & Röder, 2014). The results we report were qualitatively similar when accuracy was dichotomized with a more strict criterion of 20°, see Figure S2 in the supplemental results). We furthermore dichotomized movement speed into slow and fast by collapsing over the five slowest and five fastest movement speeds. This analysis step greatly reduces the computational demands of model fitting, but preserves the well-known modulation of higher performance during slow as compared to fast speeds under parallel instructions. Note, that we illustrate all 10 speed levels in our figures of the raw data, both for comparison with previous studies, and to demonstrate consistency across lower and higher speed levels. Finally, we subsumed hand postures into a two-leveled factor by pooling both hands down and both hands up as ‘same hand orientation’ and left up/right down and left down/right up as ‘different hand orientation’ (Heed & Röder, 2014).

In response to the concern of a reviewer that based on several earlier reports (Buchanan, Kelso, DeGuzman, & Ding, 1997; Kelso, Buchanan, DeGuzman, & Ding, 1993), we furthermore ascertained that changes in the right-left movements that we report here were not due to a transfer of movement into another movement dimension (such as up-down). We ascertained that (1) the number of movement cycles identified at each speed were comparable across speeds; (2) that the highest velocities were observed in the relevant, and not in an irrelevant, dimension; and (3) that the standard deviation of movement velocity was, accordingly, highest in the relevant dimension (see Figures S3-S6 in the supplemental results for illustration).

We fitted a hierarchical Bayesian logistic regression model to the dichotomized performance measure to estimate the probability of moving correctly in a given movement cycle through the linear combination of group-level regression beta weights and participant-level intercepts. Regression beta weights are denoted β_instruction_ for the main effect of the factor movement instruction, β_mirror_ for the main effect of the factor mirror view, β_posture_ for the main effect of the factor hand posture, and β_speed_ for the main effect of the factor speed. Furthermore, regression beta weights were included for all possible factor combinations and are denoted β_i_n_ with *i, n* denoting *i* factors interacting with *n* other factors (Liddell & Kruschke, 2014). For instance, the model parameter denoted β_instruction_mirror_posture_ represents the regression beta weight for the three-way interaction of movement instruction, mirror view, and hand posture. Beta weights were constrained to sum to zero, with the first factor level dummy-coded as 1 and the second one as -1 (β_instruction_: symmetrical=1, parallel=-1; β_mirror_: non-mirrored=1, mirrored=-1; β_posture_: same=1, different=-1; β_speed_: fast=1, slow=-1). Uninformative priors were chosen for all model parameters. Specifically, priors were modeled as normal distributions centered on zero, corresponding to a .5 probability of moving correctly. Precision, that is, the width of the normal distribution, of each prior was drawn from an inverse gamma distribution with shape parameter 1 and scale parameter .01 to allow for a large range of possible values (Gill, 2010). We re-sampled our model with several alternative specifications for uninformative priors to ensure that posterior distributions were robust. For instance, we drew the normal distributions’ precision from the inverse gamma function with shape parameter .01 and scale parameter .01, rendering qualitatively identical results (not reported).

We used JAGS version 4.0.0 (Plummer, 2015), R version 3.2.2 (R Core Team, 2016), and the R package runjags version 2.0.2-8 (Denwood, in press) to perform MCMC sampling. Specifically, we sampled 60,000 representative credible values from the joint posterior distribution of the model parameters in four independent chains.

The chains were burned in (1500 samples) and every 20th sample was saved, rendering a total of 12,000 recorded samples. Stable and accurate representation of the parameter posterior distributions was ensured visually using trace, autocorrelation, and density plots, as well as numerically by examining the effective sample size (ESS), and the shrink factor (Brooks & Gelman, 1998). All model parameters of interest had a minimum EES of 11,550, ensuring stable and accurate estimates of the limits comprising 95% of the posterior samples (i.e., their highest density interval (HDI); Kruschke, 2015).

For statistical inference, the model parameters of interest are the normalized group-level regression beta weights, which indicate the influence of each factor or factor combination (i.e., interaction) in determining the probability of moving correctly in the finger oscillation task. If the HDI of a beta weight representing a specific factor or interaction does not span zero, this implies that the factor contributes to the prediction of movement accuracy. In contrast, a HDI that spans zero indicates that a beta weight representing a specific factor does not contribute to the prediction of movement accuracy. In analogy to post-hoc testing in frequentist approaches, we assessed condition differences only if the HDI of the corresponding beta weight representing the overall effect or interaction did not span zero. For such comparisons, we contrasted the posterior predictive distributions of the factor level combinations that represented our hypotheses in the model. When multiple beta weights containing the hypothesis-relevant factors did not span zero, we took the beta weight representing the highest order interaction as the basis for whether a contrast should be evaluated or not. Contrasts are reported in the form of ^difference^a_b ^with *a, b* indicating *a* factor^ levels interacting with *b* other factor levels (Liddell & Kruschke, 2014). The distribution resulting from contrasting factor-level posterior predictive distributions are denoted as credible difference distributions. Similar to the inferential strategy applied to the beta weight posterior distributions, an HDI of a credible difference distribution that does not span zero indicates that the model predictions for the two conditions of interest are different from each other, whereas an HDI of a credible difference distribution that spans zero indicates that the model predictions for the two conditions do not differ statistically.

In the text, tables, and figures, beta weight and credible difference distributions are characterized by their mean and their upper and lower 95% HDI limit. Figures were prepared using the R package ggplot2 version 2.0.0 (Wickham, 2009).

